# Identification of a new antiphage system in *Mycobacterium* phage Butters

**DOI:** 10.1101/2023.01.03.522681

**Authors:** Hamidu T. Mohammed, Catherine Mageeney, Vassie C. Ware

## Abstract

During lysogeny temperate phages establish a truce with the bacterial host. In this state, the phage genome (prophage) is maintained within the host environment. Consequently, many prophages have evolved systems to protect the host from heterotypic viral attack. This phenomenon of prophages mediating defense of their host against competitor phages is widespread among temperate mycobacteriophages. We previously showed that the *Mycobacterium* phage Butters prophage encodes a two-component system (gp30/31) that inhibits infection from a subset of mycobacteriophages that include PurpleHaze, but not Island3. Here we show that Butters gp57r is both necessary and sufficient to inhibit infection by Island3 and other phages. Gp57r acts post-DNA injection and its antagonism results in the impairment of Island3 DNA amplification. Gp57r inhibition of Island3 is absolute with no defense escape mutants. However, mutations mapping to minor tail proteins allow PurpleHaze to overcome gp57r defense. Gp57r has a HEPN domain which is present in many proteins involved in inter-genomic conflicts, suggesting that gp57r may inhibit heterotypic phage infections via its HEPN domain. We also show that Butters gp57r has orthologues in clinical isolates of *Mycobacterium abscessus sp*. including the phage therapy candidate strain GD91 which was found to be resistant to the panel of phages tested. It is conceivable that this GD91 orthologue of gp57r may mediate resistance to the subset of phages tested. Challenges of this nature underscore the importance of elucidating mechanisms of antiphage systems and mutations that allow for escape from inhibition.

**IMPORTANCE:** The evolutionary arms race between phages and their bacteria host is ancient. During lysogeny, temperate phages establish a ceasefire with the host where they do not kill the host but derive shelter from it. Within the phenomenon of prophage-mediated defense, some temperate phages contribute genes that make their host more fit and resistant to infections by other phages. This arrangement has significance for both phage and bacterial evolutionary dynamics. Further, the prevalence of such antiphage systems poses a challenge to phage therapy. Thus, studies aimed at elucidating antiphage systems will further our understanding of phage-bacteria evolution as well as help with efforts to engineer therapeutic phages that circumvent antiphage systems.

## INTRODUCTION

Bacteriophages exert strong selective pressure on bacteria populations. Consequently, bacteria have evolved defense mechanisms that antagonize phage infections. These defense mechanisms include BREX(1), Argonaute(2), restriction modification(3), abortive infection systems(4) and the adaptive CRISPR/cas system(5) (see reviews (6, 7)). Antiphage systems that block infection of rival phages have emerged as a result of competition between phages infecting a common host. When temperate phages lysogenize the host, some prophages deploy antiphage defenses that protect the lysogen from heterotypic viral attack. Such a framework is beneficial to both host bacteria and prophage, as the bacterium is protected from heterotypic viral attack while the prophage derives shelter and maintenance of its genome(8). It is therefore not surprising that most bacterial genomes contain prophages(9, 10). Antiphage defense systems, whether bacteria-encoded or prophage-mediated, may target every step of the phage infection process(11–13).

Mycobacteriophages (a subset of the ubiquitous phage world) constitute, arguably, the largest collection of individual phages that has been isolated and characterized(14) (phagesdb.org). Comparative genome analyses reveal that mycobacteriophage genomes show great genetic diversity modulated by extensive mosaicism(15–17). Exploration of mycobacteriophages has generated a host of genetic tools used to study *Mycobacteria*(18–21). Furthermore, mycobacteriophages have been used for therapeutic purposes(22, 23). Collectively, greater insights into host-phage interactions have been uncovered from recent investigations into mycobacteriophage biology.

Prophage-mediated defense against heterotypic phage attack is pervasive among mycobacteriophages. Several types of defense mechanisms have been described. Subcluster I2 phage Sbash defends its lysogen against heterotypic phage Crossroads (subcluster L2) via an abortive infection process analogous to the Rex A/B system of *E. coli* bacteriophage Lambda(24, 25). Among cluster N temperate phages, Panchino encodes a restriction modification system (gp28) that inhibits heterotypic phage infection while Charlie gp32 is predicted to block viral attack by blocking DNA injection(26). The Phrann prophage encodes a (p)ppGpp synthetase, which upon activation, promotes the synthesis of (p)ppGpp leading to arrest of cell growth that stalls phage lytic growth(26, 27). Remarkably, all cluster N phage defenses previously characterized map to a small highly variable, centrally-located genomic region typically flanked on the left by the lysis cassette and on the right by the immunity cassette(26). Genes within this central variable region (CVR) are expressed from the prophage during lysogeny(26). Mycobacteriophage Butters (cluster N) defends its *Mycobacterium smegmatis* mc^2^155 lysogen against several heterotypic phages(26, 28). Butters has a genome size of 41,491bp, a GC content of 66%, and an overall architecture similar to other cluster N phages(29). We previously showed that Butters genes *30* and *31* define a two-component system that inhibits infection from heterotypic phages PurpleHaze (subcluster A3), and Alma (subcluster A9), but not Island3 (subcluster I1)(28). Therefore, a distinct Butters prophage-mediated mechanism, independent of genes *30* and *31*, must be considered to account for defense against Island3 infection.

Here, we report that Butters gene *57r* (on the reverse strand of the annotated forward gene *57*), is both necessary and sufficient to defend against Island3 and a host of other heterotypic phages. Using bacteriophage recombineering of electroporated DNA (BRED)(20), phage plating efficiency assays, RNAseq, and qPCR, we show that the gene product (gp) of Butters gene *57r* acts post-DNA injection and antagonizes DNA amplification of Island3. We note that a Butters gene *57r* orthologue is present in several clinical mycobacterial isolates, including *Mycobacterium abscessus*, supsp. bolletii (GD91) which showed resistance to phage infections during screening of phages for phage therapy(30). Conceivably, Butters gp57r could be instrumental in establishing resistance in several pathogenic bacterial strains against therapeutic phages, underscoring the importance of elucidating mechanisms undergirding prophage-mediated defenses.

## RESULTS

### Butters gene *57r* is both necessary and sufficient to defend against Island3

Phage plating efficiency assays show that the Butters lysogen mc^2^155(Butters) defends against Island3 by ≥ 5 orders of magnitude (5 logs)(26, 28) (Fig. 1). Given that phage defense systems characterized within prophages of cluster N mycobacteriophages have mapped to their respective CVR(26) and that phage defense systems are typically clustered in defense Islands(7, 31), we hypothesized that mc^2^155(Butters) defense against Island3 would be mediated by genes within the Butters CVR (i.e., from genes *30 to 36*).

**FIG 1.**
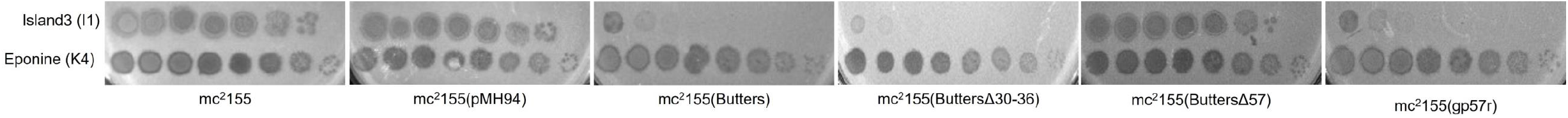
Infectivity profile of Island3 on *M. smegmatis* mc^2^155 and Butters lysogen derivatives. Phage lysates of Island3 and Eponine (used as control) were serially diluted to 10^-7^ and spotted (3 μl each) onto a lawn of each bacterium plated with 1X top agar. Island3 efficiently infects wildtype mc^2^155 and the empty vector strain [mc^2^155(pMH94)] but is inhibited on Butters lysogen [mc^2^155(Butters)] as well as mutant of Butters lysogen with genes *30-36* deleted [mc^2^155(ButtersΔ30-36)]. Mutant of Butters lysogen with gene *57* deleted [mc^2^155(ButtersΔ57)] fully restores Island3 infectivity while *M. smegmatis* mc^2^155 expressing Butters gene *57r* [mc^2^155(gp57r)] fully recapitulates inhibition of Island3 to levels comparable to that observed on Butters lysogen lawn. This panel is representative of at least three independent biological replicates for each bacteria strain, with *n>3* technical replicates. In no case did variation in EOPs between replicates exceed an order of magnitude.

To test this hypothesis, our initial strategy was to use BRED to delete the entire region containing genes *30-36* as a block and then to construct an mc^2^155(ButtersΔ30-36) lysogen from the resulting mutant phage for comparative plating efficiency assays with mc^2^155(Butters) and wild type mc^2^155. Deletion of the gene *30-36* block failed to abolish defense against Island3 (Fig. 1), indicating the likelihood that other Butters genes have a role in establishing defense. Dedrick *et al*. (2017) previously showed RNAseq patterns from a Butters lysogen that revealed broader prophage expression beyond Butters genes *30-36*, and *38* (immunity repressor), and included anti-sense expression from two regions: one spanning genes *42* and *43* and the other within gene *57*(26) (Fig. 2). Our next steps were then to explore putative roles of these genomic segments in propagating Island3 defense using BRED as a deletion strategy.

**FIG 2.**
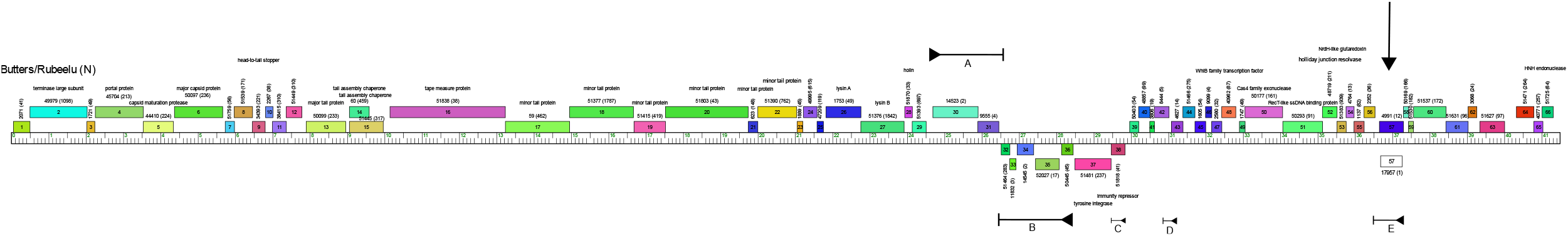
Genome map of Butters (and Rubeelu). Phamerator map(54) of Butters (and Rubeelu). Butters and Rubeelu are nearly identical except for 24 single nucleotide polymorphisms across their 41,491bp genome. Genes are depicted by colored rectangular boxes and are aligned to their genomic ruler. Gene numbers are within the rectangular boxes and putative gene functions, if known, are indicated above the gene. Numbers above the genes represent gene group names or phams and numbers in parenthesis show number of genes within the pham. Gene annotations for Butters and Rubeelu are nearly identical except for gene *57* (arrow pointing down) which is called on the forward strand in Butters (blue box) but called on the reverse strand in Rubeelu (white box). RNAseq experiment(26) show that regions A-E, denoted by a horizontal line with a shaded arrowhead (indicating direction of transcription) on one end and a short vertical line on the other end, are expressed. Thus gene *57* on the reverse strand (E) is expressed in the Butters lysogen.

Multiple attempts to delete the *42-43* region to recover mutant phages were unsuccessful, suggesting a requirement of this genomic region for a productive phage infection. Within the Butters genome, gene *57* is annotated on the forward strand but in cluster N mycobacteriophage Rubeelu (which is nearly identical to Butters except for 24 single nucleotide polymorphisms (SNPs) across their 41,491 bp genomes), gene *57* is annotated on the reverse strand (Fig. 2). Hence, we will refer to the open reading frame (ORF) on the forward strand as gene *57f* and that on the reverse strand as gene *57r*, with their corresponding gene products as gp57f and gp57r, respectively. The entire ORF of gene *57r* is contained within gene *57f* (Fig. 2). Deletion of gene *57* to yield mc^2^155(ButtersΔ57) fully restored infection by Island3 (Fig. 1). To test whether gene *57r* (the gene expressed in RNAseq) is sufficient to defend against Island3, we cloned Butters gene *57r* into the integrative proficient vector pMH94(19) and transformed it into electrocompetent *M. smegmatis* to yield mc^*2*^155(gp57r). The gene *57r* construct included 100bp of the region upstream of the annotated start, capturing the predicted native promoter and ribosome binding site. Plating efficiency assays showed that mc^2^155(gp57r) recapitulated defense against Island3 comparable to that mounted by mc^2^155(Butters) (Fig. 1).

Butters gp57r defense against Island3 appears absolute as we were unable to isolate Island3 defense escape mutants that can infect mc^2^155(gp57r) or mc^2^155(Butters). However, a few plaques form (between 5-12) following Island3 infection of mc^2^155(Butters). Analysis of these plaques shows they contain recombinant phages formed from genomes of the invading Island3 phage and the resident Butters prophage. The left and right arms of the recombinant phage genome are derived from the resident Butters prophage while the middle (which makes up ~20% of the resulting genome) is derived from Island3 (and are thereby named “BIBs”). Isolation and characterization of these novel recombinant genomes is described elsewhere (Mohammed *et al*., 2022, submitted). Plaques resulting from Island3 infection of mc^2^155(ButtersΔ57) yields predominantly wildtype Island3 phages, but about 1 in 100 are recombinants (data not shown), suggesting that recombination between the Island3 and Butters genomes is independent of gp57r defense.

### Butters gp57r defense extends to phages in other clusters

Phage sensitivity assays were expanded to determine if defense conferred by Butters gp57r was unique to the subcluster I1 phage Island3. Results show that Butters gp57r defends against several other phages (Fig. 3A). Like Island3, defense against fellow subcluster I1 phage Brujita, and the subcluster A1 phage Norz appears absolute and yields no defense escape mutants (Fig. 3A). However, phages such as LHTSCC (A4), PurpleHaze (A3), Perplexer (A4) and Bud (A11) showed partial infection that may be due to defense escape mutants (Fig. 3A, 3B). For PurpleHaze, we were able to isolate and characterize eleven defense escape mutants that resolve into 6 unique mutational events in genes encoding minor tail proteins (Table 1). These PurpleHaze DEMs efficiently infect defense-competent strains, mc^2^155(Butters) and mc^2^155(gp57r) (Fig. 3B). These mutations encompass a spectrum of mutation types, including nonsense mutations, frameshifts, readthroughs and missense mutations. These results suggest possible interactions between the gp57r defense system and minor tail proteins of some heterotypic phages.

**FIG 3.**
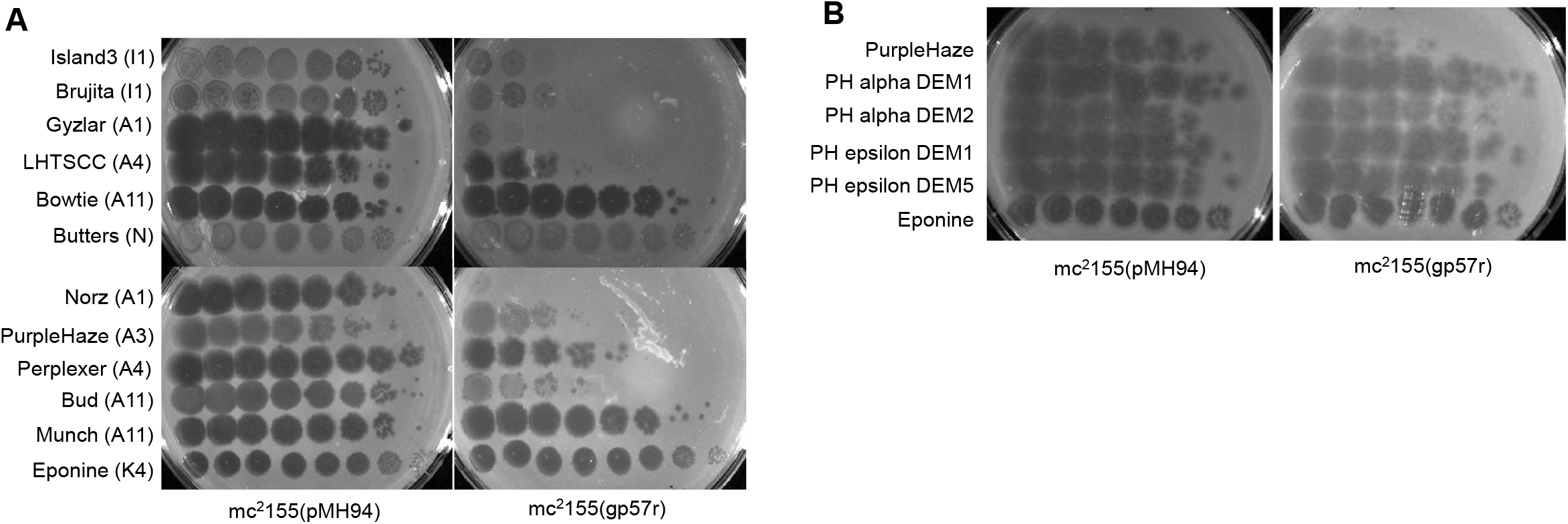
Butters gp57r defense against heterotypic phages. Plating efficiencies of heterotypic phages on *M. smegmatis* mc^2^155 strains expressing pMH94 empty vector and Butters gene *57r* [designated mc^2^155(pMH94) and mc^2^155(gp57r) respectively]. Phages spotted are listed on the left with clusters or subclusters in parenthesis. Phage lysates were serially diluted to 10^-7^ and spotted (3 μl each) onto a lawn of each bacterium plated with 1X top agar. (A) Butters gp57r inhibits infection of a subset of heterotypic phages. On mc^2^155(gp57r) lawn, the absence of individual plaques in the dilution series for Island3, Brujita, and Norz suggests that observed clearings are due to “killing from without” and not productive infection. (B) PurpleHaze defense escape mutants (PH DEMs) efficiently infect mc^2^155(gp57r). Alpha DEMs were isolated from Butters lysogen lawn [mc^2^155(Butters)] while epsilon DEMs were isolated from the mc^2^155 strain expressing only gp57r [mc^2^155(gp57r)]. Panels are representative of at least three independent biological replicates for each bacteria strain, with *n=2* technical replicates. In no case did variation in EOPs between replicates exceed an order of magnitude.

**Table 1.** Mutations within PurpleHaze defense escape mutants.

| **Mutant** | **DNA sequence changes** | **Amino acid changes** | **Remarks** |
| --- | --- | --- | --- |
| PH alpha DEM1 | g.1790T>C and<br>g.2818A>G | p.Val36Ala, and<br>p.Thr379Ala | Two missense<br>mutations |
| <sup>a</sup>PH alpha DEM2,3 | g.1606G>A | p.Thr81* | Premature stop codon |
| PH alpha DEM4 | g.2154_2161del | Several missense<br>mutations | Frameshifts,<br>premature stop |
| PH alpha DEM5,<br>epsilon DEM1,2,3 | g.2883A>G | p.*400Trp | Abolishes stop,<br>readthrough |
| PH epsilon DEM4,5 | g.2879_2880insG | N/A | Abolishes stop,<br>readthrough |
| PH alpha DEM6 | g.2001G>A | p.Trp106* | Premature stop |

PH denotes PurpleHaze. Alpha DEMs1-6 were isolated from Butters lysogen lawn [*mc*<sup>2</sup>155(Butters)] while epsilon DEMs1-5 were isolated from the *mc*<sup>2</sup>155 strain expressing inly gp57r [*mc*<sup>2</sup>155(gp57r)]. Whole genome sequences were obtained for alpha DEMs1-6. For Epsilon DEMs1-5, analysis was based on Sanger sequencing of the PCR amplification of the genomic region with coordinates 1199-3032. <sup>a</sup>DEM2 and DEM3 have mutations in PurpleHaze gene 3. All other DEMs have mutations in gene 4.

### Bioinformatics analysis of Butters gp57r reveals the presence of a HEPN domain

We used a gamut of bioinformatics tools to characterize the Butters gp57r ORF. A model of gp57r is shown in Supplementary Fig. 1A. Membrane domain prediction tool DeepTMHMM(32) reveals no transmembrane domain within Butters gp57r (supplementary Fig. 1B). HHpred analysis showed weak homology of Butters gp57r with CRISPR-associated (Cas) DxTHG family (probability = 55.77% and E-value = 170), HEPN pEK499 p136 (probability = 55.25% and E-value = 47), Mannitol repressor (probability = 40.5% and E-value = 300), and Abi-like protein (probability = 35.57% and E-value = 88)(33–35). A key observation is that these hits align with amino acids 94-99 (RDRCAH) of gp57r and show homology with an RX4-6H motif that is highly conserved in HEPN domains of proteins implicated in genomic conflicts and pathogen targeting(36) (Supplementary Fig. 1A).

An NCBI BLAST(n) search shows that gene *57r* (present in Butters and Rubeelu) is present in 11 additional mycobacteriophages. Apart from Xavia (a subcluster P3 phage), all other hits are cluster N mycobacteriophages. It should be noted that not all cluster N phages have this gene segment (neither *57f* nor *57r*). The multiple sequence alignment program Clustal Omega sorts gp57r homologues into 3 groups based on amino acid identities(37) (Supplementary Fig. 1C). Group 1 includes Butters and Rubeelu as identical members. Xavia is the only member of Group 2, and it is 84.82% identical to Butters/Rubeelu homologues. Group 3 has 10 members: nine of them (ShrimpFriedEgg, Purgamenstris. Nenae, Spongebob, BabeRuth, PhancyPhin, Snekmaggedon, Jamie19, and Redi) have identical sequences and are 80.00% identical to Butters/Rubeelu homologues. The last member of Group 3, Raymond7, is 99.47% identical to other members and 79.47% identical to the Butters homologues. All these homologues have the conserved RX4-6H motif within the HEPN domain.

Differences in gp57r amino acid composition among phages may contribute to differences in Island3 infection among cluster N lysogens. It is notable that lysogens of ShrimpFriedEgg and Redi fail to inhibit Island3 infection(26, 28). Nevertheless, transcriptional analysis of the Redi lysogen shows expression from the genetic region harboring this gene(26). Since the Redi gp57r homologue (and by extrapolation, ShrimpFriedEgg) is expressed during lysogeny, we suggest that the inability of the Redi/ShrimpFriedEgg homologue to inhibit Island3 infection may be attributable to the 20% amino acid dissimilarity between the Butters and Redi/ShrimpFriedEgg sequences (Supplementary Fig. 2). This observation will be critical in future explorations to determine regions/motifs within Butters gp57r that are required for defense against Island3 infection.

### Arrest of phage infection occurs downstream of DNA injection

Phage infection is initiated with phage adsorption to bacteria followed by phage genome injection into the host cell, expression of the phage genome, assembly of progeny phages, and bacterial lysis to release newly assembled phage. In principle, each stage of the phage infection process can be targeted by an antiviral defense system as revealed in many examples(11, 13). We performed an adsorption assay to test whether gp57r impairs Island3 adsorption. Results show that Island3 absorption is not impaired in the mc^2^155(gp57r) strain (Fig. 4A).

**FIG 4.**
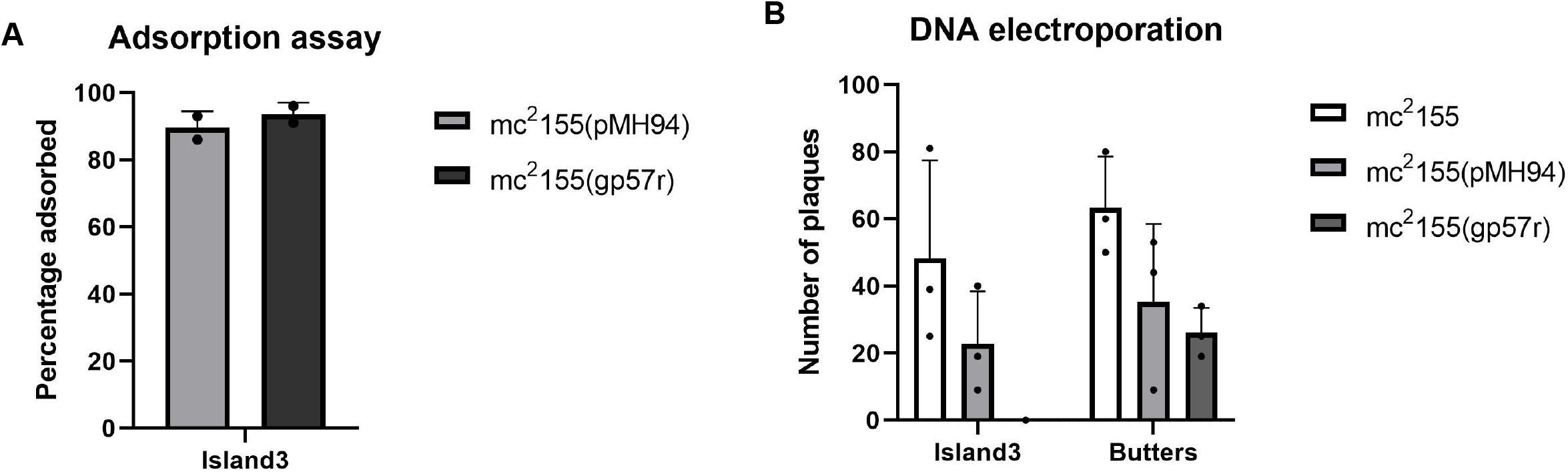
Effects of Butters gp57r on adsorption and DNA injection of Island3. (A). Butters gp57r does not impair the ability of Island3 to adsorb to mc^2^155(gp57r) cells. Experiment was performed at least 3 times each with 2 biological replicates. (B). DNA electroporation assay. Electroporation of Island3 DNA produces plaques on mc^2^155 and mc^2^155(pMH94) but not on mc^2^155(gp57r). Butters DNA serving as control produces plaques on all cell types transformed. Data represents 3 technical replicates of the same experiment.

Next, we reasoned that, if gp57r blocks Island3 DNA injection, then electroporation (a process that circumvents adsorption and DNA injection steps) of Island3 DNA into mc^2^155(gp57r) should yield a productive infection and generate viral progeny. Electroporation of Butters DNA (serving as a control) successfully infected all strains: electrocompetent mc^2^155, mc^2^155(pMH94) and mc^2^155(gp57r). In contrast, electroporation of Island3 DNA was successful only in transforming gp57r-deficient strains, mc^2^155 and mc^2^155(pMH94), yielding several plaques. Electroporation of Island3 DNA into mc^2^155(gp57) failed to transform the strain to yield plaques, despite the fact that adsorption and DNA injection were bypassed (Fig. 4B). These results favor the proposal that gp57r acts downstream of DNA injection.

### Island3 nucleic acid metabolism is disrupted in defense-competent strains

Comprehensive analysis of HEPN domain-containing proteins shows that most function as RNases, especially proteins with the conserved RX4-6H motif(36). Given the presence of a HEPN domain and the conserved RX4-6H motif within Butters gp57r, we explored the hypothesis that Butters gp57r antagonizes Island3 infection by targeting specific Island3 transcripts. We performed RNAseq and differential gene analyses to quantify Island3 transcripts generated during early and late infection in mc^2^155(Butters), mc^2^155(gp57r), and wildtype mc^2^155 (control).

Consistent with the transcription profiles of other mycobacteriophages with similar genomic architecture(26), Island3 genome transcription starts from a rightwards promoter upstream of gene *35* (Fig. 5A). During early infection of mc^2^155 (i.e., 30 mins post infection [mpi]), we observe modest expression of genes *35-62* and then expression tapers off (Fig. 5A). Although functions of several genes within this region are unknown, many others are predicted to play various roles in DNA maintenance. Proposed functions in this genomic region include: a helix-turn-helix (HTH) protein as the putative excisionase (gene *35*), an antirepressor that promotes lytic gene expression(38) (gene *36*), the RecE/RecT system that mediates homologous recombination(39) (genes *48,49*), resolvase (gene *53*), and a DNA methylase (gene *55*). By late infection (150 mpi), DNA maintenance gene expression remains modest, but we observe robust expression levels from genes *69*-*76* and from genes *1-32*. (Fig. 5A). Only gene *75* (n-acetylglucosaminyltransferase) has a known function within the gene *69-76* block. Genes *1-32* encode viral structural and morphogenetic genes such as terminase (gene *2*), portal protein (gene *4*), capsid maturation protease (gene *6*), major capsid protein (gene *7*), major tail protein (gene *13*), tape measure protein (gene *16*), minor tail proteins (genes *17, 19-21*) and genes required for cell lysis: lysin proteins (genes *29,30*) and a putative holin (gene *31*) (Fig. 5A).

**FIG 5.**
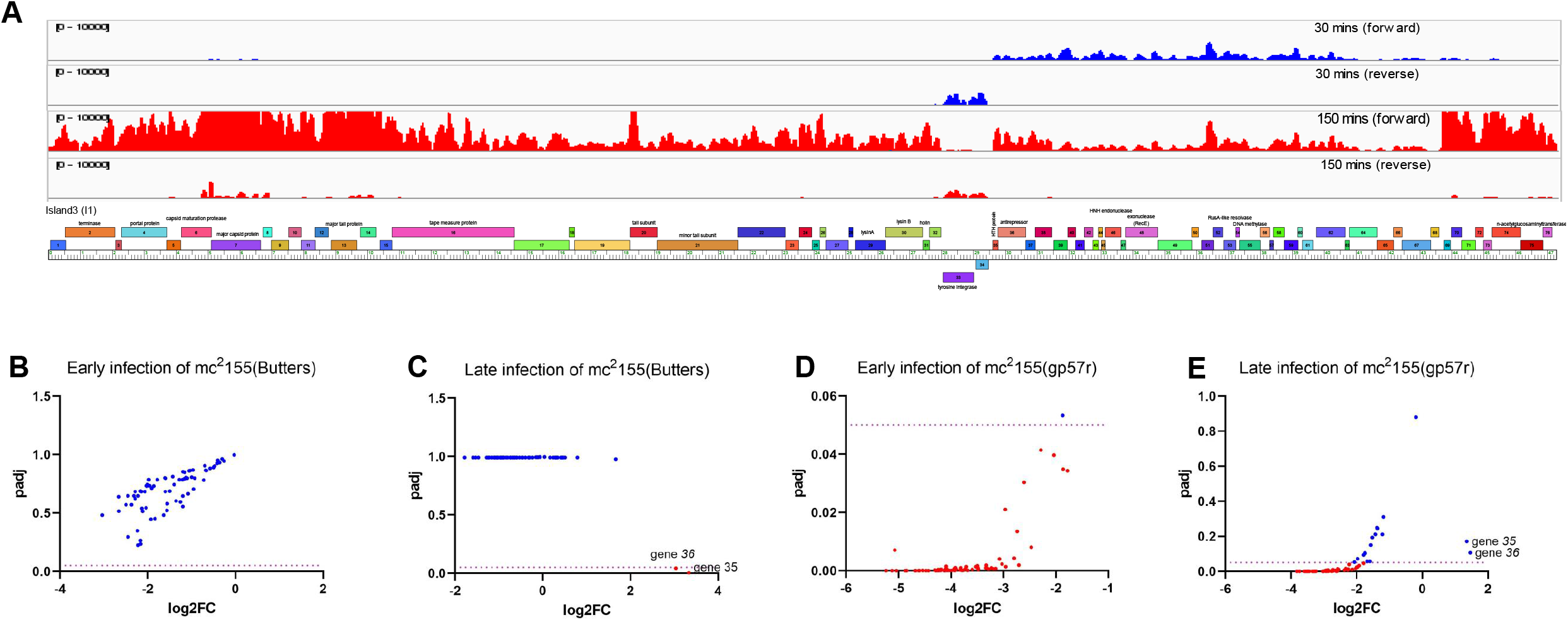
RNAseq analysis of Island3. RNA was isolated at 30 mpi and 150 mpi, representing early and late infection respectively, during Island3 infection of mc^2^155, mc^2^155(Butters) and mc^2^155(gp57r). (A) RNAseq profile of Island3 during infection of mc^2^155. Map showing coverage of Island3 transcripts on both forward (blue) and reverse (red) DNA strands at the indicated time points. Coverage tracks generated with IGV(53) and aligned to Island3 genome map generated with Phamerator(54). (B-E) Differential gene expression analysis of RNAseq data during Island3 infection of (B) mc^2^155(Butters) at 30 mpi, (C) mc^2^155(Butters) at 150 mpi, (D) mc^2^155(gp57r) at 30 mpi, (E) mc^2^155(gp57r) at 150 mpi.

Given that the Butters lysogen defends against Island3 infection, we examined the transcription profile of Island3 during infection of mc^2^155(Butters) to determine if there were notable differences relative to Island3 infection of mc^2^155.

During early infection of mc^2^155(Butters), no Island3 gene is differentially expressed (padj > 0.05), although transcript levels are generally lower compared to infection of the control (mc^2^155) (Fig. 5B). However, during late infection, genes *35* (HTH protein), and *36* (antirepressor) are upregulated (padj < 0.05) and no other Island3 gene is differentially expressed (Fig. 5C). Next, we analyzed the transcription profile of Island3 during infection of mc^2^155(gp57r) since this strain recapitulated inhibition of Island3 infection as in the lysogen (Fig. 1). Here, differences in gene expression are notable at both early and late time points when compared to infection of the control (mc^2^155). During early infection of mc^2^155(gp57r) strain, all Island3 transcripts except gene *30* are significantly downregulated with fold changes ranging from 3-38X (Fig. 5D). At 150 mpi, Island3 genes *1-22*, and *39-76* are downregulated with fold change differences between 3-14X, but genes *23-38* are not differentially expressed (padj > 0.05) (Fig. 5E).

Remarkably, genes *35 and 36* appear upregulated although padj fails to meet the threshold of significance (Fig. 5E). The parallels from analyses of Island3 transcripts during infection of defense competent strains [mc^2^155(Butters) and mc^2^155(gp57r)] are that there are generally lower transcript levels during early phases of infection and elevated levels of genes *35 and 36* during late infection (Figs 5B–5E). Collectively, these data do not reveal any unique Island3 transcript candidates that may be targets for specific degradation in the mc^2^155(Butters) or mc^2^155(gp57r) strains. Given that all Island3 transcripts are made, including those encoded by structural assembly genes, it is noteworthy that not a single Island3 plaque was observed during plaque assays. It is possible gp57r antagonism of Island3 infection may occur elsewhere in the infection process with generalized downregulation of transcription being an outcome of another primary gp57r event.

Next, we tested whether the Island3 genome is amplified or not. First, a timed phage infection assay was established to gauge a time point where the greatest probability of phage DNA amplification or virion particle assembly would occur prior to cell lysis. The Island3 phage titer from infected mc^2^155(pMH94) remained fairly constant from time point T = 0 to T = 2.5 hours but increased approximately 10-fold at T = 3.5 hours presumably after lysis and phage release into the medium (Fig. 6A). Based on this time course, Island3 DNA amplification was compared at T = 2.5 hours during infection of mc^2^155(gp57r) to infection of the mc^2^155(pMH94) control. Quantitative PCR (qPCR) was used to estimate the relative genome amplification of Island3 during infection of mc^2^155(gp57r) compared to infection of the mc^2^155(pMH94) control. At T = 2.5 hours after infection, we observed a 25% increase in Island3 genome quantity in the mc^2^155(pMH94) control strain, but a 38% decrease in the amount of Island3 genome in the mc^2^155(gp57r) strain (Fig. 6B). Within the experimental set up that allows for bacterial growth, we would expect a relative fold change increase in the amount of the Island3 genome if its amplification exceeds bacterial genome amplification but a relative fold change decrease if phage genome amplification remains unchanged. Thus, our results suggest that the Island3 genome is not amplified during infection of the mc^2^155(gp57r) strain.

**FIG 6.**
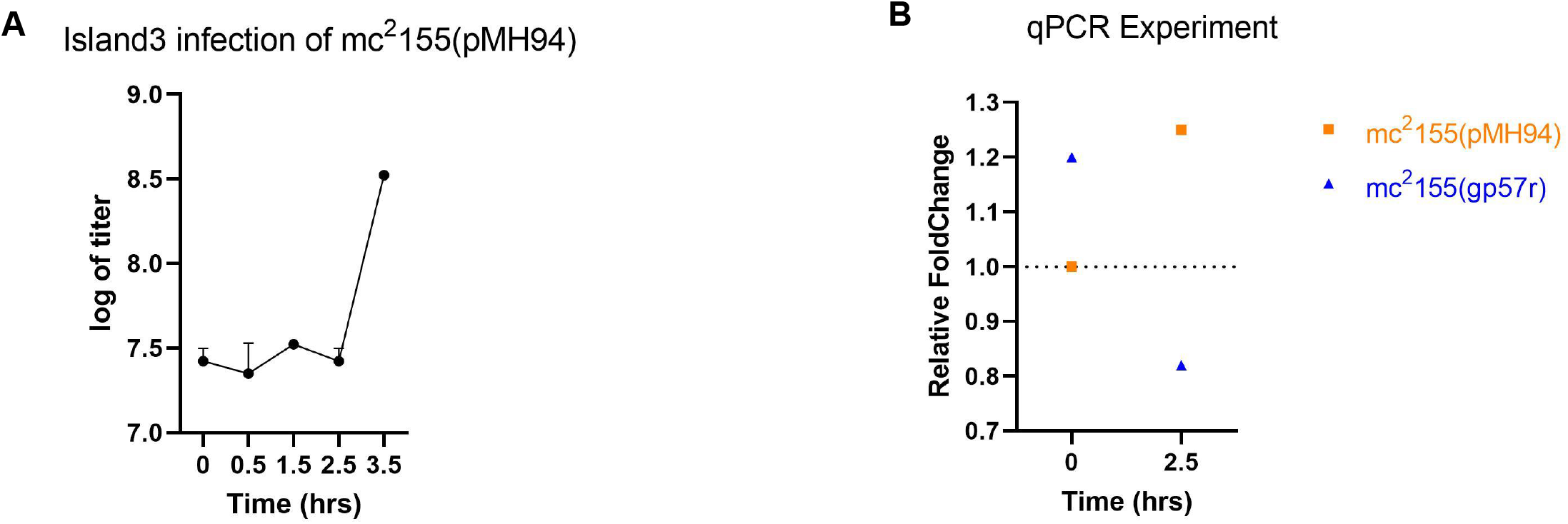
Dynamics of Island3 genome amplification during infection. (A). Time course for Island3 infection of mc^2^155(pMH94). We set up Island3 infection of mc^2^155(pMH94) at MOI = 0.1. Samples were drawn at the indicated time points for titer determination using spot test assay. (B). Measuring Island3 genome amplification by relative qPCR. The delta-delta Ct method was used to calculate the fold change in Island3 genome normalized to infection of mc^2^155(pMH94) control at T0. This result is representative of 3 independent experiments. Each experiment was set up on two biological replicates and 3 technical replicates. We observed variation in fold change at T0 but irrespective of the quantity at T0, at T2.5, Island3 genome quantity increases in mc^2^155(pMH94) but decreases in mc^2^155(gp57r).

## DISCUSSION

Bacteria are under constant bombardment by phage infection. Consequently, bacteria have evolved a plethora of defense systems against phage attack with phages evolving counter defense strategies(6, 26). Beyond building armament against bacterial hosts, phages also compete with other phages infecting a common host. Thus, over the course of their evolution, some phages have evolved defense systems that target other competitor phages. While some phages deploy antiphage defenses during lytic infection(40, 41), the majority of phage antagonism against other phages plays out during lysogeny where there is the greatest opportunity for an invading phage to encounter a resident prophage. Because such prophage-mediated defenses ultimately benefit the bacterial host, it is unsurprising that prophages abound in many bacterial genomes(9, 10). Predictably, several prophage-host collusions against viral attack have been discovered(12, 25, 26, 28, 42). Elucidating prophage-mediated defense mechanisms is crucial to understanding biological interactions between bacteria and their viruses within various ecological systems. Moreover, such knowledge could be critical in engineering therapeutic phages that evade defense systems.

The prophage of mycobacteriophage Butters mediates defense against heterotypic phage attack. Our earlier work showed that Butters genes *30* and *31* defend against a select panel of phages, but not Island3. Given the preponderance of antiviral genes to cluster within defense islands and the fact that all characterized cluster N prophage defense systems cluster within their CVRs, we focused on genes within the CVR as likely candidates that function in defense against Island3 infection. As shown in Fig. 1, deletion of the entire Butters CVR (*30-36*) failed to abolish defense against island3. Deletion of gene *57*, however, abolished defense against Island3 (Fig. 1). Phage sensitivity assays on mc^2^155(gp57r) recapitulated defense against Island3 to an equivalent extent as defense observed when Island3 is plated on a wildtype Butters lysogen (Fig. 1). Thus, Butters gp57r is both necessary and sufficient to defend against Island3.

Collectively, three lines of evidence support the hypothesis that Butters gp57r exerts its antiviral activity within the cytoplasm. First, bioinformatics analyses using the transmembrane domain prediction tool TMHMM predict no transmembrane domains within gp57r. Next, phage adsorption assays show no impairment in Island3 adsorption in strains expressing gp57r (mc^2^155[gp57r]) compared to the control (mc^2^155[pMH94]). Finally, electroporation (a process that bypasses phage adsorption and DNA injection) of Island3 DNA into the mc^2^155(gp57r) strain still results in inhibition of Island3 infection (no plaques are recovered) while parallel electroporation of DNA from a phage not subject to gp57r inhibition or into control strains yields plaques. In all, evidence favors an interpretation of an intracellular mechanism for gp57r antagonism of Island3 infection.

Bioinformatic analysis shows that gp57r includes a HEPN domain, recognized as a signature domain for proteins typically acting as RNases that are involved in intergenomic conflicts(36). RNAseq analyses showed overexpression of only two Island3 genes (*35* and *36*) during late infection of mc^2^155(Butters). During late infection of mc^2^155(gp57r), some Island3 genes were downregulated between 3-14-fold levels. All transcripts are present and yet no Island3 phages are released from infection of mc^2^155(Butters). RNAseq analysis did not identify any specific transcript as a potential target of gp57r antagonism, but it remains unclear what threshold levels of expression for any specific transcripts are required for a productive infection. Thus, it is unknown if gp57r antagonism involves targeting of specific transcripts for degradation possibly through an HEPN domain.

In the absence of clear evidence for specific targeting of Island3 transcripts, additional mechanisms to account for inhibition of Island3 infection should be considered. In principle, the inhibitory activity of gp57r could work to inhibit translation of Island3 transcripts, inhibit Island3 DNA amplification, or even block cellular lysis as has been demonstrated in a *Bacillus subtilis* system(43). Yet given the penchant of HEPN domains to antagonize nucleic acids, we investigated the possibility that Island3 DNA amplification is impaired within the mc^2^155(gp57r) strain. Quantitative PCR results suggest that little, if any, Island3 DNA is made during Island3 infection of mc^2^155(gp57r). This constraint appears to be a key reason accounting for lack of viral progeny during infection of the mc^2^155(gp57r) strain. Diminished DNA amplification may also account for the observed downregulation of Island3 transcripts since newly amplified DNA serves as templates for robust transcription and is subsequently packaged into new virions.

The gp57r defense system described here features several characteristics described for the abortive infection system mediated by the AbiK of *Lactococcus lactis*. Like Butters gp57r, AbiK is a standalone system that contains a HEPN domain(36). AbiK abolishes infection by P335 by inhibiting ERF/Rad52-like ss annealing protein of P335, resulting in abolition of P335 DNA amplification(44–46). Although the ERF/Rad52 family of proteins generally functions in homologous recombination(39, 47), this group has also been implicated in repair of dsDNA breaks(48), and in directing the formation of the primosome required for DNA replication(49). The Rad52 functional homologue in Island3 is gene *49* (putative RecT) which is essential for its lytic cycle (our unpublished data). One possibility is that Butters gp57r antagonizes Island3 RecT and inhibits recombinase function which abrogates DNA replication. Alternatively, gp57r may function analogously as the HEPN domain-containing PrrC and RloC which inhibit enterobacteriophage T4 infection by digesting tRNA^Lys^ and abolishing translation(50). Inhibiting translation of proteins required for DNA replication could produce phenotypes consistent with gp57r results. Further, gp57r could act as a toxin kept in an inactive state until activation by interaction with a phage component leading to cell death. This mechanism has been described for several other HEPN domain-containing Abi systems(36).

Gp57r is constitutively expressed from the Butters prophage even in the absence of a phage challenge. Its expression in *M. smegmatis* is neither bactericidal nor bacteriostatic, as no differences in growth are observed for the mc^2^155(gp57r) strain compared to the control mc^2^155 strain (our unpublished data). We propose a model to account for Island3 inhibition by Butters gp57r where adsorption and DNA injection of the invading Island3 DNA proceeds normally. At a point in late gene expression (150 mpi), newly synthesized phage components or metabolic pressure derived from the infection process activates gp57r to exert its inhibitory effect. For PurpleHaze, phages that escape defense mounted by gp57r and produce infectious particles have loss-of-function mutations that map to minor tail protein genes *3* and *4*. Thus, PurpleHaze minor tail proteins PH_3 and PH_4 are likely components that activate gp57r directly or indirectly through other protein interactions. While minor tail proteins can be essential during the early stages of infection (e.g., during adsorption and DNA injection), it is unknown if production of minor tails during late infection could serve as targets for antiphage defense systems. That PurpleHaze defense escape mutants with minor tail protein defects remain competent for infection suggests a degree of structural redundancy among minor tail components in building an infectious PurpleHaze particle. Unlike PurpleHaze, no Island3 defense escape mutants were recovered. Whatever the nature of the Island3 component that activates gp57r activity, it would likely be essential for phage production to account for the lack of infectious mutant particles. While the outcome of gp57r antagonism against Island3 results in impairment of DNA amplification, defense against PurpleHaze and other phages may proceed through different mechanisms.

Overexpression of Island3 genes *35* (putative excise) and *36* (antirepressor) at 150 mpi during infection of defense-competent strains is rather enigmatic. It is unknown what the impact of overexpression of these genes has on host metabolism. But, given the canonical roles of these two proteins to drive the phage into the lytic cycle, overexpression of these genes may be a strategy to accomplish this. Yet, Island3 does not proceed into an efficient lytic cycle to produce phages in the presence of gp57r.

Phage resistance systems, whether bacteria-encoded or prophage-mediated have been predicted to present key challenges to phage therapy(51,52). *Mycobacterium* phages have been used to treat *Mycobacteria abscessus* infections (22, 23). Butters gene *57r* is found in several *M. abscessus* strains isolated in human clinical samples (Table 2) and its antiphage activities could pose challenges to phage therapy. For example, *M. abscessus subsp. bolletii* strain GD91 was found among the S morphotype strains that are resistant to phage infection during phage susceptibility screening to identify phages for phage therapy(30). Strain GD91 carries a prophage that encodes an orthologue of Butters gp57r (Table 2). It is conceivable that this GD91 orthologue of gp57r may mediate resistance to a subset of phages that were tested. Challenges of this nature necessitate research into characterization of antiphage systems and mutations that allow for escape from inhibition. Thus, the importance of elucidating mechanisms undergirding antiphage systems cannot be overemphasized.

**Table 2.**
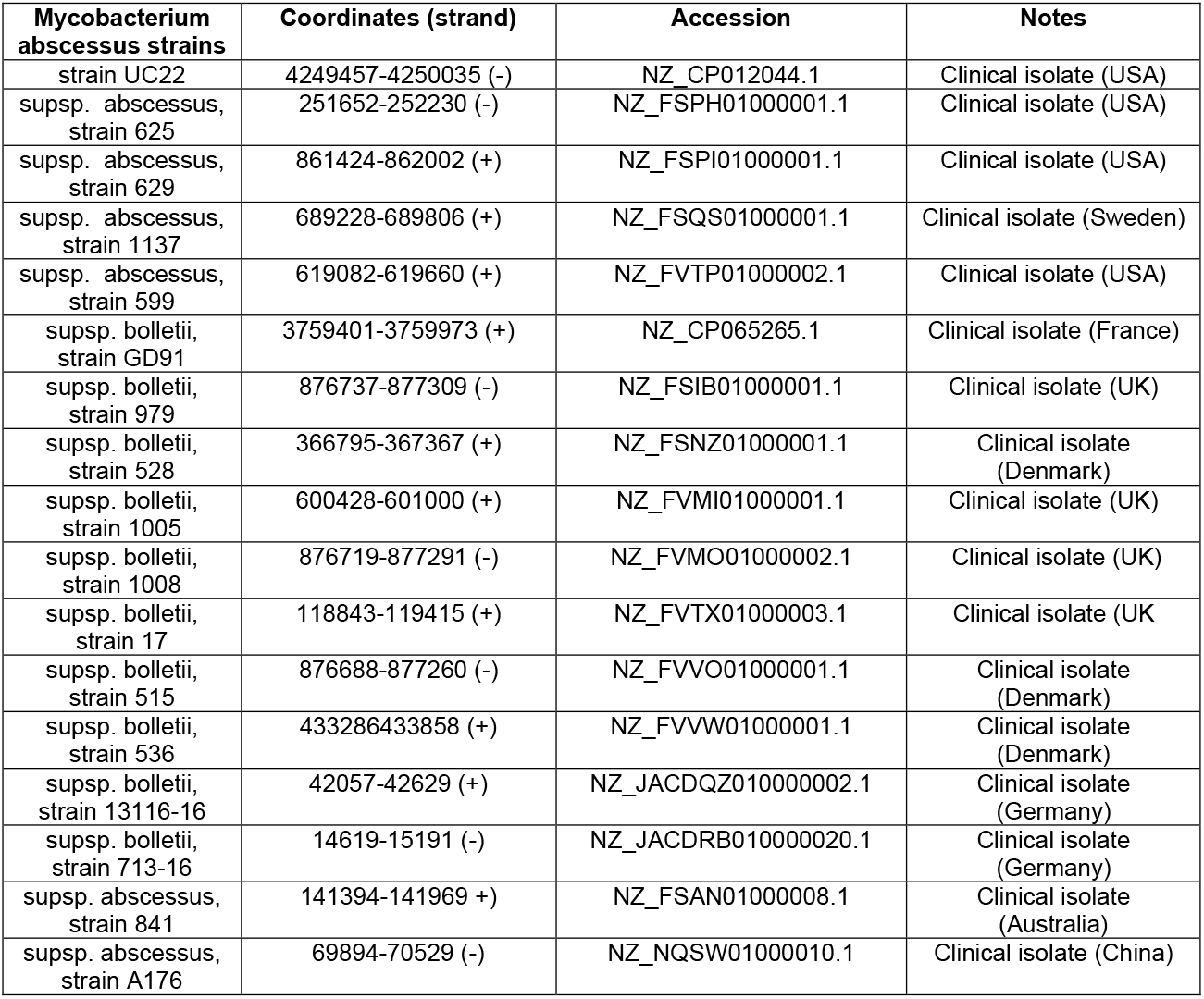
Clinical isolates of *Mycobacterium abscessus* strains with a Butters gp57r orthologue.

## METHODS

### Phage isolation, propagation, and genomic analysis

Phages [GenBank accession numbers] used in this study are Butters [**KC576783**], Island3 [**HM152765**], Eponine [**MN945904**], Brujita [**FJ168659**], Gyzlar [**MZ355726**], LHTSCC [**JN699015**], Bowtie [**MK524511**], Norz [**ON970624**], PurpleHaze [**KY965063**], Perplexer [**OP910130**], Bud [**MK524514**] and Munch [**MK524513**]. Phages were isolated and grown on *Mycobacterium smegmatis* mc^2^155 as described in (29). Island3, Brujita, LHTSCC, and PurpleHaze lysates was obtained from the Hatfull lab (University of Pittsburgh). The genomic sequence for the Island3 strain used in this study differs from that of the wild type with a 257-bp deletion (coordinates 43307 to 43563) and a C2656T SNP. Phage lysates (titers, ≥1 ×10^9^ PFU/ml) were diluted with phage buffer (0.01 M Tris, pH 7.5, 0.01 M MgSO4, 0.068 M NaCl, and 1 mM CaCl_2_). Whole genome sequences exist for all phages used in this study except PurpleHaze epsilon DEMs. For these, we used PCR to amplify the genomic region from within gene *2* to within gene *5* (coordinates 1199-3032). PCR products were purified using PCR cleanup kit (Promega). Sanger sequencing was performed at Psomagen, Inc.

### Construction of mc^2^155(ButtersΔ57) strain

The BRED technique(20) was used to delete the entire ORF of Butters gene *57r* (spanning coordinates 36651-37292) except its last 4 bp overlap with gene *58*. We generated two PCR products, a 450 bp product upstream (PCR1) and a 183 bp downstream (PCR2) of the deleted segment, respectively. Primers were designed to yield a 50 bp overlap between the two PCR products [See detailed strategy in (28)]. The equal amounts of PCR1 and PCR2 were used as template for BRED substrate (PCR3) using forward primer from PCR1 and reverse primer from PCR2. The resulting PCR-generated substrate was used for BRED after agarose gel purification, PCR cleanup (Promega), and quantification. Purified substrate (200 ng) and 200 ng of Butters genomic DNA were co-electroporated into recombineering-efficient strain *M. smegmatis* mc^2^155 carrying plasmid pJV53. Cell recovery, plating, PCR screening, plaque purification, and amplification were conducted as previously described(20). Primers used are in Supplementary Table 1. Q5 High-Fidelity 2X Master Mix (New England Biolabs) was used for all PCR reactions.

### Construction of mc^2^155(gp57r) strain

Recombinant strain to express Butters gene *57r* was created as follows. All primers used in this study are shown in Table S1. Butters genome segment (coordinates 36681-37356) was cloned into the XbaI site of integration proficient, kanamycin (KAN)-resistant, and ampicillin (AMP)-resistant vector pMH94(19) using conventional restriction enzyme/ligation methods. This gene segment contained all the ORF of gene *57r* and 100 bp of the upstream sequences containing the endogenous promoter and ribosome binding site. PCR primers (Integrated DNA Technologies) were designed with XbaI site at the ends. Phage genes were amplified from Butters genomic DNA by PCR using Q5 high-fidelity DNA polymerase (New England Biolabs). PCR products were digested with XbaI overnight (O/N), purified by gel extraction, and ligated into XbaI-digested pMH94 using T4 DNA ligase (New England Biolabs) at 16°C O/N. Chemically competent *E. coli* were transformed and plated onto Kan/Amp plates, and colonies were screened by PCR with primers flanking the cloning site. Recombinant plasmids were verified by sequencing (Psomagen). Electrocompetent *M. smegmatis* mc^2^155 cells were prepared and transformed with recombinant pMH94 plasmids. After recovery, cells were plated on selective medium containing Luria broth agar with 50_μg/ml kanamycin. Strains were grown in 7H9 complete medium [.e. 7H9 medium enriched with albumin (5%) and dextrose (2%) (AD supplement), 1 mM CaCl_2_, 50_μg/ml kanamycin, 50_μg/ml carbenicillin (CB), and 10_μg/ml cycloheximide (CHX)] for 5 days at 37°C.

### Phage Sensitivity Assays

Bacteria lawns were made by plating 250 μl of the bacteria culture with 3.5 ml of 1X top agar (at ~55°C) on an LB agar plate (with 10 μg/ml of cycloheximide [CHX] and 50 μg/ml of carbenicillin [CB]). Phage lysates were serially diluted to 10^-7^ and spotted (3μl each) onto lawns of interest. Plates were incubated for 48 h at 37°C. Phage growth was assessed at 24 and 48 h. EOP was determined as titer on experimental strain/titer on *M. smegmatis* mc^2^155.

### Phage adsorption assay

An aliquot of a high titer lysate of Island3 was added to 1 ml of appropriate culture [mc^2^155(pMH94) or mc^2^155(gp57r)] to yield MOI = 0.1. The mixture was incubated for 15 mins at 37°C with shaking to allow for adsorption of phage particles. After incubation, samples were spun at 10,000 rpm for 1 min to pellet phagebacteria complexes. Resulting supernatants were serially diluted and spotted on mc^2^155 lawns to determine the titer of unadsorbed phages. This titer was subtracted from initial phage concentration (formed on adding phage to bacteria) to yield concentration of adsorbed phages. Fraction of adsorbed phages were determined and converted to percentages. The experiment was performed 3 times and two biological replicates were used on each occasion.

### DNA electroporation assay

Phage DNA (200 ng) isolated using phenol:chloroform method was electroporated into 100 μl of electrocompetent mc^2^155, mc^2^155(pMH94), and mc^2^155(gp57r) respectively. Cells were recovered in 1 ml of 7H9 complete medium for 2 hours at 37°C with shaking. Cells were pelleted at 10,000 rpm for 1 min and resuspended with 200 μl of media. Cells were added to 250 μl of mc^2^155 and plated with 3.5ml of 1X top agar unto LB agar plate (with 10 μg/ml of cycloheximide [CHX] and 50 μg/ml of carbenicillin [CB]. Plaques were counted after 48 hours of incubation at 37°C.

### Time course assay

Island3 infection of mc^2^155(pMH94) was set up at MOI = 0.1. Samples were taken at the indicated time points, serially diluted and titered using a spot test assay.

### RNAseq analysis

Island3 infection of bacterial cultures were set up at MOI = 1 and incubated at 37°C with shaking. Samples were at taken 30 mins and at 150 mins, pelleted at 10,000 rpm for 1 min and flash frozen until RNA isolation. The RNA preparation, library preparation, and RNA-seq analysis were done at Azenta/Genewiz, USA. Total RNA was extracted from fresh frozen cell pellets using Qiagen RNeasy Plus Universal mini kit following manufacturer’s instructions (Qiagen, Hilden, Germany). Extracted RNA samples were treated with TURBO DNase (Thermo Fisher Scientific, Waltham, MA, USA) to remove DNA following manufacturer’s protocol. RNA samples were quantified using Qubit 2.0 Fluorometer (Life Technologies, Carlsbad, CA, USA) and RNA integrity was checked using Agilent TapeStation 4200 (Agilent Technologies, Palo Alto, CA, USA). rRNA depletion sequencing library was prepared by using QIAGEN Fastselect rRNA 5S/16S/23S Kit (Qiagen, Hilden, Germany). Strand-specific RNA sequencing library was prepared by using NEBNext Ultra II Directional RNA Library Prep Kit for Illumina following manufacturer’s instructions (NEB, Ipswich, MA, USA). Briefly, the enriched RNAs were fragmented for 8 minutes at 94 °C. First strand and second strand cDNA were subsequently synthesized. The second strand of cDNA was marked by incorporating dUTP during the synthesis. cDNA fragments were adenylated at 3’ends, and indexed adapter was ligated to cDNA fragments. Limited cycle PCR was used for library enrichment. The incorporated dUTP in second strand cDNA quenched the amplification of second strand, which helped to preserve the strand specificity. The sequencing library was validated on the Agilent TapeStation (Agilent Technologies, Palo Alto, CA, USA), and quantified by using Qubit 2.0 Fluorometer (ThermoFisher Scientific, Waltham, MA, USA) as well as by quantitative PCR (KAPA Biosystems, Wilmington, MA, USA). The sequencing libraries were multiplexed and clustered on a flowcell. After clustering, the flowcell was loaded on the Illumina instrument (HiSeq 4000 or equivalent) according to manufacturer’s instructions. The samples were sequenced using a 2×150bp Paired End (PE) configuration. Image analysis and base calling were conducted by the Control software. Raw sequence data (.bcl files) generated from the sequencer were converted into fastq files and demultiplexed using Illumina’s bcl2fastq 2.17 software. One mismatch was allowed for index sequence identification. After demultiplexing, sequence data were checked for overall quality and yield. Then, sequence reads were trimmed to remove possible adapter sequences and nucleotides with poor quality using Trimmomatic v.0.36. The trimmed reads were mapped to the reference genome using the STAR aligner v.2.5.2b. BAM files were generated as a result of this step. Unique gene hit counts were calculated by using featureCounts from the Subread package v.1.5.2 Only unique reads within exon regions were counted. After extraction of gene hit counts, the gene hit counts table was used for downstream differential expression analysis. Using DESeq2, a comparison of gene expression between the groups of samples was performed. The Wald test was used to generate p-values and Log2 fold changes. Genes with adjusted p-values < 0.05 and absolute log2 fold changes > 1 were called as differentially expressed genes for each comparison. Integrative Genomic Viewer(53) was used to visualize bam files produced from alignment of reads to reference genomes.

### Quantitative PCR experiment and primer design

Island3 infection of 1 ml of mc^2^155(pMH94) and mc^2^155(gp57r) were set up at MOI = 0.1 and incubated for 10 mins at 37°C with shaking. Afterwards, cells were pelleted and resuspended with 0.4% 1N HCl and incubated for 1 min to inactivate residual unadsorbed phages. Cells were pelleted and washed 3X using 7H9 complete and finally suspended in 1 ml 7H9 complete media and incubated at 37°C with shaking. Samples were taken after the washes (designated T0) and 150 mins (designated T2.5) post infection for qPCR. DNA was isolated by boiling100 μl of samples for qPCR for 5 mins and pelleted at 10,000 rpm for 1 min. Supernatant was collected, and DNA concentration estimated with a nanodrop. qPCR was set up on two biological replicates and three technical replicates using <u>Applied Biosystems™</u> PowerTrack <u>™ SYBR Green Master Mix</u>. qPCR was performed using Applied Biosystems 7300 Real-Time PCR System. Primers (Supplemental Table 1) designed to amplify a segment of Island3 DNA (target) and mc^2^155 (control) were ordered from Integrated DNA Technologies. The delta-delta Ct method was used to calculate the fold change in Island3 genome quantity.

### Data availability

The genome sequences of all phages used in this study are available at https://phagesdb.org. GenBank accession numbers are provided in Materials and Methods. Constructs and sequences from this study are available upon request. RNAseq data sets have been deposited in the Gene Expression Omnibus (GEO) under accession no. GSE221406.

## SUPPLEMENTARY DATA

**FIG S1**

**FIG S2 Table S1**

## ACKNOWLEDGEMENTS

We thank SEA-PHAGES team members Daniel Russell and Rebecca Garlena for performing genome sequencing, genome assembly, and deposition of raw reads into the Sequence Read Archive. We thank the Graham Hatfull laboratory for Island3, Brujita, LHTSCC, and PurpleHaze phage lysates, Ariana Amendolara for helping with primer design, members of the Ware lab, Dr. Michael Kuchka and the phage community for thoughtful feedback about this work.

---

**Table S1:**
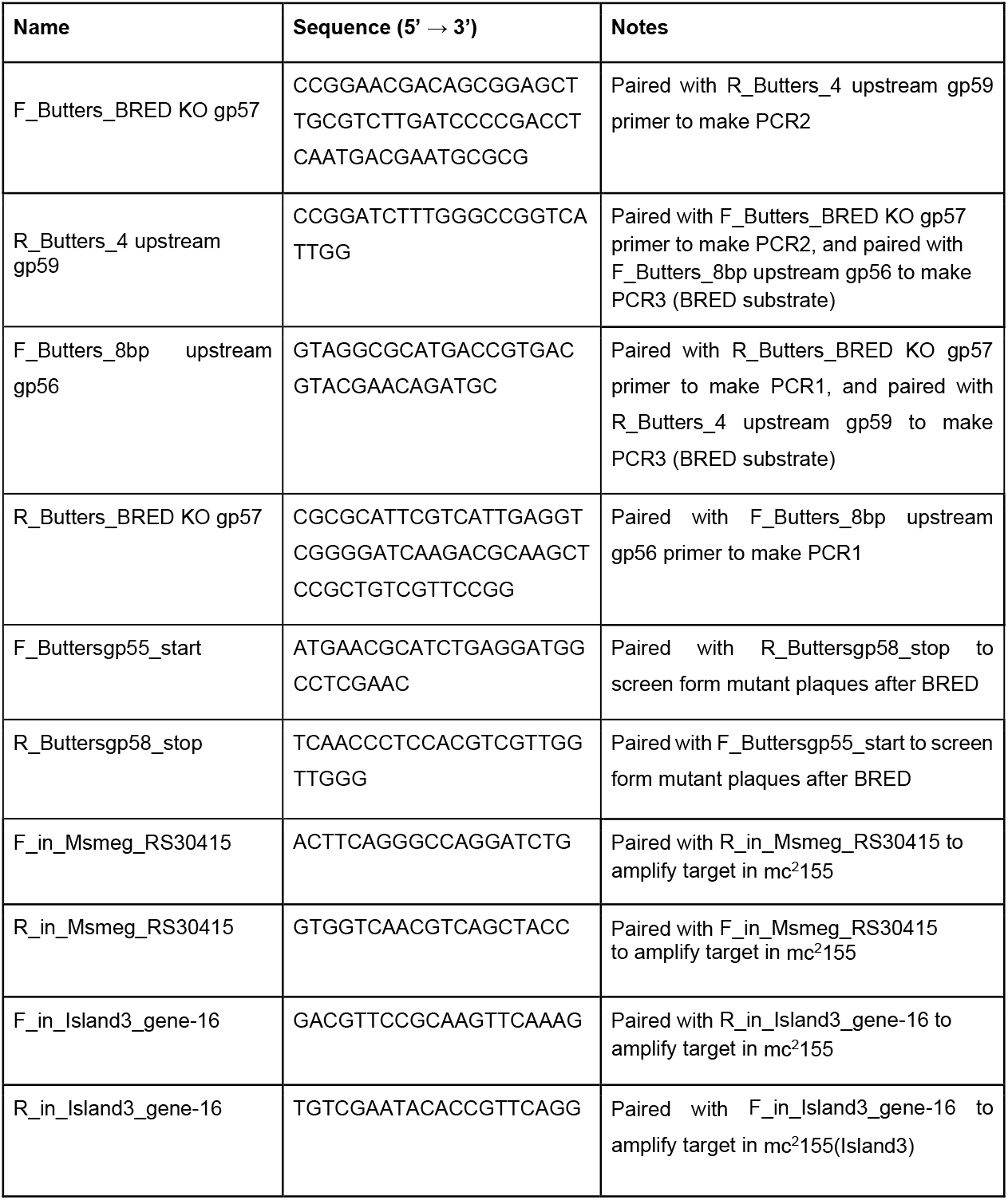
List of primers used in this study.

**Supplementary Fig. S1.**
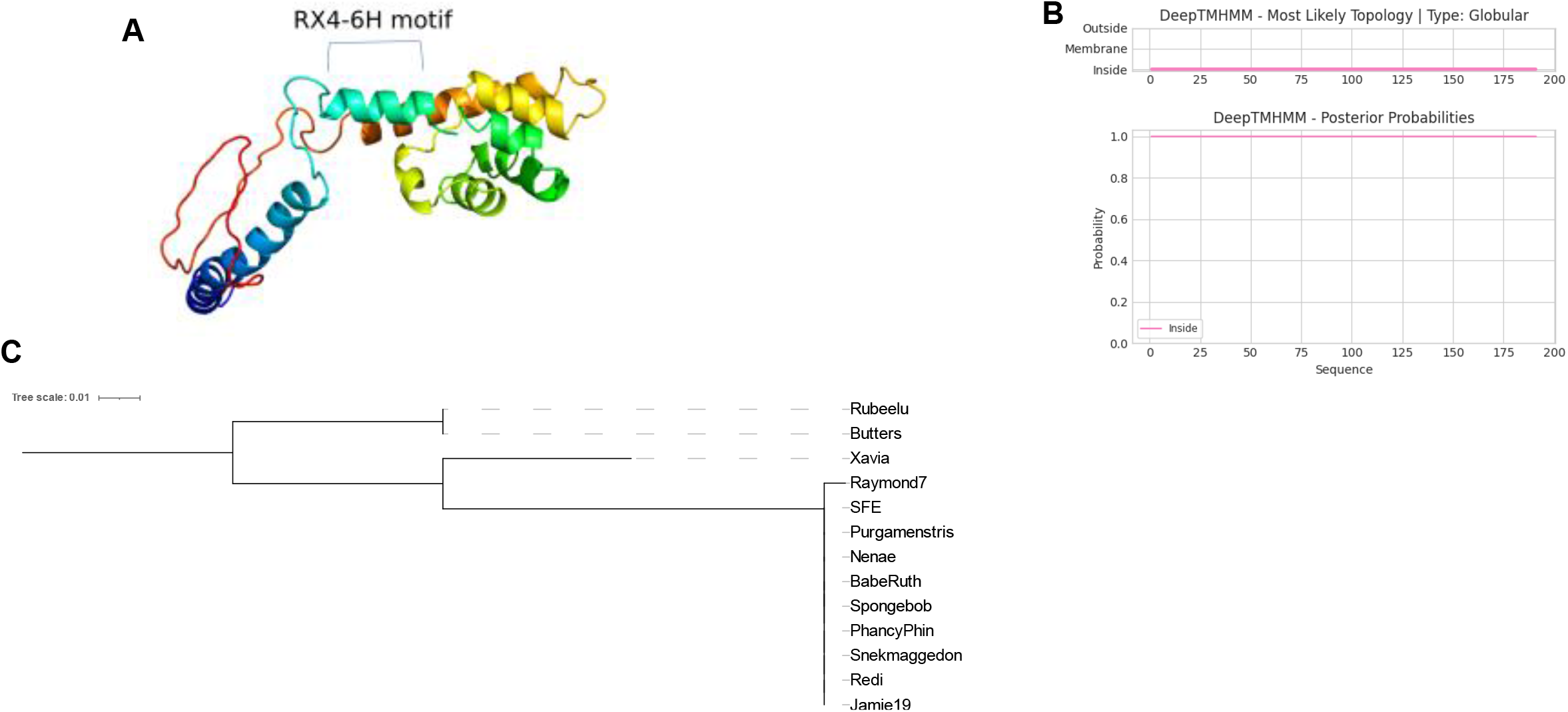
Bioinformatic analyses of Butters gp57r. (A) Predicted secondary structure of gp57r. A phyre2(1) model of gp57r. The helix containing the conserved RX4-6H motif is indicated. (B)) DeepTMHMM(2) prediction of membrane topology. The amino acid index is shown on the horizontal axis. Gp57r has no predicted transmembrane. (C) Phylogeny of gp57r. Clustal Omega(3) was used to analyze phylogenetic relationship between homologues of gp57r. iTOL(4) was used to view the resulting tree.

**Supplementary Fig. S2.**
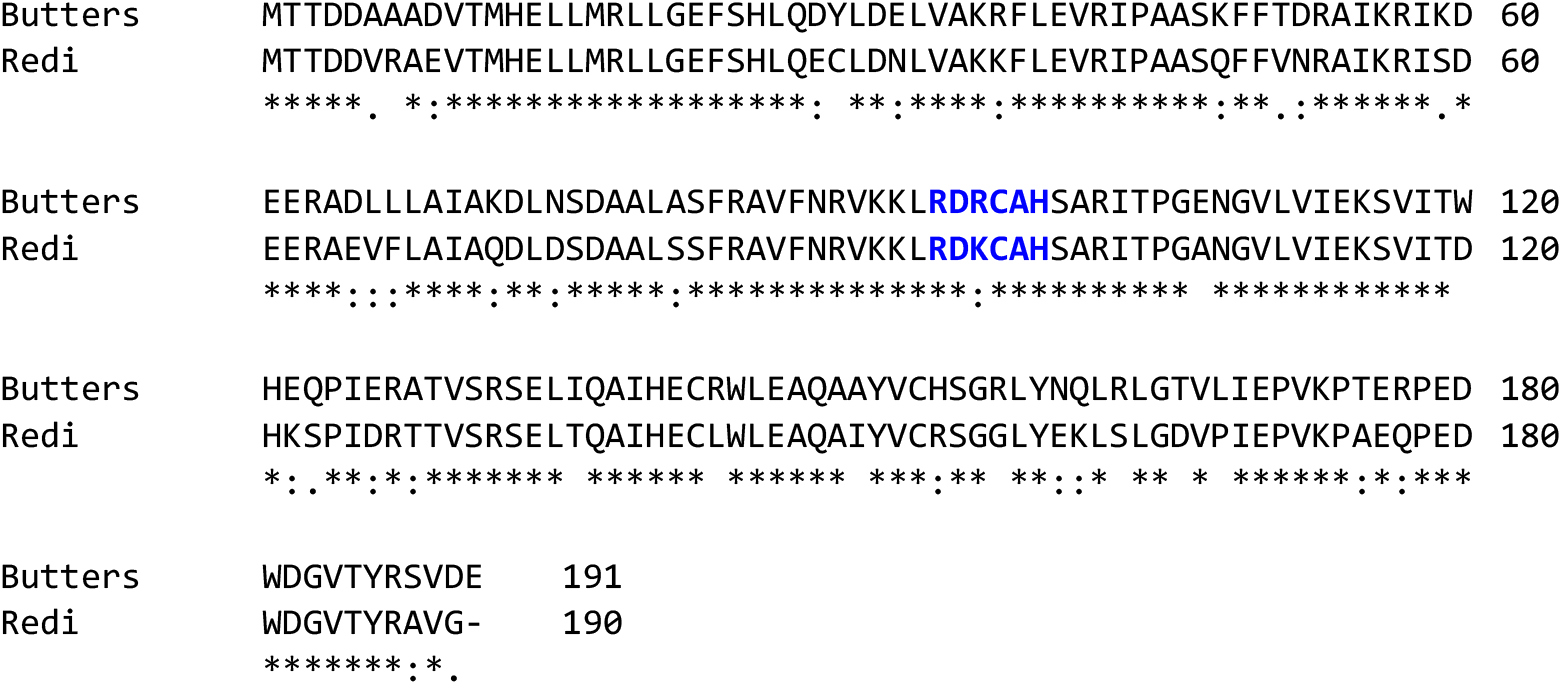
Sequence alignment of Butters gp57r and its Redi homologue. Clustal Omega(5) was used to align amino acid sequences of Butters gp57r and its Redi homologue. [*] denotes a position that is fully conserved, [:] denotes conservation of amino acid with strongly similar properties and [.] denotes conservation of amino acid with weakly similar properties. The RDRCAH/RDKCAH highlighted (blue and bold font) is a conserved motif within the HEPN domain of proteins implicated in pathogen targeting(6). While both homologues are expressed in their respective lysogens, Island3 infection is inhibited by only the Butters lysogen (but not Redi lysogen)(7, 8). This difference in defense capabilities may be due to the amino acid dissimilarity shown here.

## REFERENCES

1. Goldfarb T, Sberro H, Weinstock E, Cohen O, Doron S, Charpak-Amikam Y, Afik S, Ofir G, Sorek R. 2015. BREX is a novel phage resistance system widespread in microbial genomes. EMBO J 34:169–183.

2. Swarts DC, Jore MM, Westra ER, Zhu Y, Janssen JH, Snijders AP, Wang Y, Patel DJ, Berenguer J, Brouns SJJ, Van Der Oost J. 2014. DNA-guided DNA interference by a prokaryotic Argonaute. Nature 507:258–261.

3. Tock MR, Dryden DTF. 2005. The biology of restriction and anti-restriction. Curr Opin Microbiol 8:466–472.

4. Chopin MC, Chopin A, Bidnenko E. 2005. Phage abortive infection in lactococci: Variations on a theme. Curr Opin Microbiol 8:473–479.

5. Barrangou R, Fremaux C, Deveau H, Richards M, Boyaval P, Moineau S, Romero DA, Horvath P. 2007. CRISPR provides acquired resistance against viruses in prokaryotes. Science (80-) 315.

6. Hampton HG, Watson BNJ, Fineran PC. 2020. The arms race between bacteria and their phage foes. Nature. Nature Research.

7. Makarova KS, Wolf YI, Koonin E V. 2013. Comparative genomics of defense systems in archaea and bacteria. Nucleic Acids Res 41:4360–4377.

8. Feiner R, Argov T, Rabinovich L, Sigal N, Borovok I, Herskovits AA. 2015. A new perspective on lysogeny: Prophages as active regulatory switches of bacteria. Nat Rev Microbiol. Nature Publishing Group.

9. Casjens S. 2003. Prophages and bacterial genomics: What have we learned so far? Mol Microbiol.

10. Canchaya C, Proux C, Fournous G, Bruttin A, Brüssow H. 2003. Prophage Genomics. Microbiol Mol Biol Rev 67:238–276.

11. Labrie SJ, Samson JE, Moineau S. 2010. Bacteriophage resistance mechanisms. Nat Rev Microbiol.

12. Bondy-Denomy J, Qian J, Westra ER, Buckling A, Guttman DS, Davidson AR, Maxwell KL. 2016. Prophages mediate defense against phage infection through diverse mechanisms. ISME J 10:2854–2866.

13. Egido JE, Costa AR, Aparicio-Maldonado C, Haas P-J, Brouns SJJ. 2021. Mechanisms and clinical importance of bacteriophage resistance. FEMS Microbiol Rev https://doi.org/10.1093/femsre/fuab048.

14. Hatfull GF. 2022. Mycobacteriophages: From Petri dish to patient. PLoS Pathog 18.

15. Pedulla ML, Ford ME, Houtz JM, Karthikeyan T, Wadsworth C, Lewis JA, Jacobs-Sera D, Falbo J, Gross J, Pannunzio NR, Brucker W, Kumar V, Kandasamy J, Keenan L, Bardarov S, Kriakov J, Lawrence JG, Jacobs WR, Hendrix RW, Hatfull GF. 2003. Origins of highly mosaic mycobacteriophage genomes. Cell 113:171–182.

16. Jacobs-Sera D, Marinelli LJ, Bowman C, Broussard GW, Guerrero Bustamante C, Boyle MM, Petrova ZO, Dedrick RM, Pope WH, Modlin RL, Hendrix RW, Hatfull GF. 2012. On the nature of mycobacteriophage diversity and host preference. Virology.

17. Pope WH, Bowman CA, Russell DA, Jacobs-Sera D, Asai DJ, Cresawn SG, Jacobs WR, Hendrix RW, Lawrence JG, Hatfull GF. 2015. Whole genome comparison of a large collection of mycobacteriophages reveals a continuum of phage genetic diversity. Elife 4.

18. Jacobs WR, Tuckman M, Bloom BR. 1987. Introduction of foreign DNA into mycobacteria using a shuttle phasmid. Nature 327:532–535.

19. Hong Lee M, PASCOPELLAt L, Jacobs WR, Hatfull GF. 1991. Site-specific integration of mycobacteriophage L5: Integration-proficient vectors for Mycobacterium smegmatis, Mycobacterium tuberculosis, and bacille Calmette-Guerin (pathogenesis/Mycobacterium leprae/site-specific recombination/vaccines)Proc. Natl. Acad. Sci. USA.

20. Marinelli LJ, Piuri M, Swigoňová Z, Balachandran A, Oldfield LM, van Kessel JC, Hatfull GF. 2008. BRED: A simple and powerful tool for constructing mutant and recombinant bacteriophage genomes. PLoS One 3.

21. Van Kessel JC, Hatfull GF. 2008. Mycobacterial recombineering. Methods Mol Biol 435.

22. Rebekah M. Dedrick, Bailey E. Smith, Madison Cristinziano, Krista G. Freeman, Deborah Jacobs-Sera, Yvonne Belessis, A. Whitney Brown, Keira A. Cohen, Rebecca M. Davidson, David van Duin, Andrew Gainey, Cristina Berastegui Garcia, C.R. Robert George, Ghady GFH. 2022. Phage Therapy of Mycobacterium Infections: Compassionate-use of Phages in Twenty Patients with Drug-Resistant Mycobacterial Disease. Clin Infect Dis 1–6.

23. Dedrick RM, Guerrero-Bustamante CA, Garlena RA, Russell DA, Ford K, Harris K, Gilmour KC, Soothill J, Jacobs-Sera D, Schooley RT, Hatfull GF, Spencer H. 2019. Engineered bacteriophages for treatment of a patient with a disseminated drug-resistant Mycobacterium abscessus. Nat Med 25:730–733.

24. Parma DH, Snyder M, Sobolevski S, Nawroz M, Brody E, Gold L. 1992. The Rex system of bacteriophage λ: Tolerance and altruistic cell death. Genes Dev 6:497–510.

25. Gentile GM, Wetzel KS, Dedrick RM, Montgomery MT, Garlena RA, Jacobs-Sera D, Hatfull GF. 2019. More evidence of collusion: A new prophage-mediated viral defense system encoded by mycobacteriophage sbash. MBio 10:1–20.

26. Dedrick RM, Jacobs-Sera D, Guerrero Bustamante CA, Garlena RA, Mavrich TN, Pope WH, Cervantes Reyes JC, Russell DA, Adair T, Alvey R, Bonilla JA, Bricker JS, Brown BR, Byrnes D, Cresawn SG, Davis WB, Dickson LA, Edgington NP, Findley AM, Golebiewska U, Grose JH, Hayes CF, Hughes LE, Hutchison KW, Isern S, Johnson AA, Kenna MA, Klyczek KK, Mageeney CM, Michael SF, Molloy SD, Montgomery MT, Neitzel J, Page ST, Pizzorno MC, Poxleitner MK, Rinehart CA, Robinson CJ, Rubin MR, Teyim JN, Vazquez E, Ware VC, Washington J, Hatfull GF. 2017. Prophage-mediated defence against viral attack and viral counter-defence. Nat Microbiol 2.

27. Potrykus K, Cashel M. 2008. (p)ppGpp: Still magical? Annu Rev Microbiol.

28. Mageeney CM, Mohammed HT, Dies M, Anbari S, Cudkevich N, Chen Y, Buceta J, Ware VC. 2020. *Mycobacterium* Phage Butters-Encoded Proteins Contribute to Host Defense against Viral Attack. mSystems 5.

29. Caratenuto RA, Ciabattoni GO, DesGranges NJ, Drost CL, Gao L, Gipson B, Kahler NC, Kirven NA, Melehani JC, Patel K, Rokes AB, Seth RA, West MC, Alhout AA, Akoto FF, Capogna N, Cudkevich N, Graham LH, Grapel MS, Haleem MM, Korenberg JB, Lichak BP, McKinley LN, Mendello KR, Murphy CE, Pyfer LM, Ramirez WA, Reisner JR, Swope RH, Thoonkuzhy MJ, Vargas LA, Veliz CA, Volpe KR, Zhang KD, Faltine-Gonzalez DZ, Zuilkoski CM, Mageeney CM, Mohammed HT, Kenna MA, Ware VC. 2019. Genome sequences of six cluster n mycobacteriophages, kevin1, nenae, parmesanjohn, shrimpfriedegg, smurph, and spongebob, isolated on mycobacterium smegmatis mc^2^155. Microbiol Resour Announc 8.

30. Dedrick RM, Smith BE, Garlena RA, Russell DA, Aull HG, Mahalingam V, Divens AM, Guerrero-Bustamante CA, Zack KM, Abad L, Gauthier CH, Jacobs-Sera D, Hatfull GF. 2021. Mycobacterium abscessus strain morphotype determines phage susceptibility, the repertoire of therapeutically useful phages, and phage resistance. MBio 12.

31. Doron S, Melamed S, Ofir G, Leavitt A, Lopatina A, Keren M, Amitai G, Sorek R. 2018. Systematic discovery of antiphage defense systems in the microbial pangenome. Science (80-) 359.

32. Hallgren J, Tsirigos KD, Damgaard Pedersen M, Juan J, Armenteros A, Marcatili P, Nielsen H, Krogh A, Winther O. 2022. DeepTMHMM predicts alpha and beta transmembrane proteins using deep neural networks https://doi.org/10.1101/2022.04.08.487609.

33. Söding J. 2005. Protein homology detection by HMM-HMM comparison. Bioinforma Orig Pap 21:951–960.

34. Zimmermann L, Stephens A, Nam SZ, Rau D, Kübler J, Lozajic M, Gabler F, Söding J, Lupas AN, Alva V. 2018. A Completely Reimplemented MPI Bioinformatics Toolkit with a New HHpred Server at its Core. J Mol Biol 430:2237–2243.

35. Gabler F, Nam S-Z, Till S, Mirdita M, Steinegger M, Söding J, Lupas AN, Alva V. 2020. Protein sequence analysis using the MPI bioinformatics toolkit. Curr Protoc Bioinforma e108 72:108.

36. Anantharaman V, Makarova KS, Burroughs AM, Koonin E V., Aravind L. 2013. Comprehensive analysis of the HEPN superfamily: Identification of novel roles in intra-genomic conflicts, defense, pathogenesis and RNA processing. Biol Direct 8.

37. Madeira F, Pearce M, Tivey ARN, Basutkar P, Lee J, Edbali O, Madhusoodanan N, Kolesnikov A, Lopez R. 2022. Search and sequence analysis tools services from EMBL-EBI in 2022. Nucleic Acids Res 50.

38. Kim M, Ryu S. 2013. Antirepression System Associated with the Life Cycle Switch in the Temperate Podoviridae Phage SPC32H. J Virol 87.

39. De Paepe M, Hutinet G, Son O, Amarir-Bouhram J, Schbath S, Petit MA. 2014. Temperate Phages Acquire DNA from Defective Prophages by Relaxed Homologous Recombination: The Role of Rad52-Like Recombinases. PLoS Genet 10.

40. Lu MJ, Henning U. 1994. Superinfection exclusion by T-even-type coliphages. Trends Microbiol 2:137–139.

41. Ko CC, Hatfull GF. 2018. Mycobacteriophage Fruitloop gp52 inactivates Wag31 (DivIVA) to prevent heterotypic superinfection. Mol Microbiol 108:443–460.

42. Montgomery MT, Guerrero Bustamante CA, Dedrick RM, Jacobs-Sera D, Hatfull GF. 2019. Yet more evidence of collusion: A new viral defense system encoded by gordonia phage carolann. MBio 10.

43. Guo L, Sattler L, Shafqat S, Graumann PL, Bramkamp M. 2022. A Bacterial Dynamin-Like Protein Confers a Novel Phage Resistance Strategy on the Population Level in Bacillus subtilis. MBio 13.

44. Emond E, Holler BJ, Boucher I, Vandenbergh PA, Vedamuthu ER, Kondo JK, Moineau S. 1997. Phenotypic and genetic characterization of the bacteriophage abortive infection mechanism AbiK from Lactococcus lactis. Appl Environ Microbiol 63:1274–1283.

45. Boucher I, Émond E, Dion E, Montpetit D, Moineau S. 2000. Microbiological and molecular impacts of AbiK on the lytic cycle of Lactococcus lactis phages of the 936 and P335 species. Microbiology 146:445–453.

46. Bouchard JD, Moineau S. 2004. Lactococcal phage genes involved in sensitivity to AbiK and their relation to single-strand annealing proteins. J Bacteriol 186.

47. Lopes A, Amarir-Bouhram J, Faure G, Petit MA, Guerois R. 2010. Detection of novel recombinases in bacteriophage genomes unveils Rad52, Rad51 and Gp2.5 remote homologs. Nucleic Acids Res 38.

48. Kolodner R, Hall SD, Luisi-DeLuca C. 1994. Homologous pairing proteins encoded by the Escherichia coii recE and recT genes. Mol Microbiol.

49. Ayora S, Missich R, Mesa P, Lurz R, Yang S, Egelman EH, Alonso JC. 2002. Homologous-pairing activity of the Bacillus subtilis bacteriophage SPP1 replication protein G35P. J Biol Chem 277.

50. Uzan M, Miller ES. 2010. Post-transcriptional control by bacteriophage T4: mRNA decay and inhibition of translation initiation. Virol J 7:1–22.

51. Caflisch KM, Suh GA, Patel R. 2019. Biological challenges of phage therapy and proposed solutions: a literature review. Expert Rev Anti Infect Ther.

52. Azam AH, Tanji Y. 2019. Bacteriophage-host arm race: an update on the mechanism of phage resistance in bacteria and revenge of the phage with the perspective for phage therapy. Appl Microbiol Biotechnol.

53. Thorvaldsdóttir H, Robinson JT, Mesirov JP. 2013. Integrative Genomics Viewer (IGV): High-performance genomics data visualization and exploration. Brief Bioinform 14.

54. Cresawn SG, Bogel M, Day N, Jacobs-Sera D, Hendrix RW, Hatfull GF. 2011. Phamerator: A bioinformatic tool for comparative bacteriophage genomics. BMC Bioinformatics 12.

## REFERENCES

1. Kelley LA, Mezulis S, Yates CM, Wass MN, Sternberg MJE. 2015. The Phyre2 web portal for protein modeling, prediction and analysis. Nat Protoc 10:845–858.

2. Hallgren J, Tsirigos KD, Damgaard Pedersen M, Juan J, Armenteros A, Marcatili P, Nielsen H, Krogh A, Winther O. 2022. DeepTMHMM predicts alpha and beta transmembrane proteins using deep neural networks https://doi.org/10.1101/2022.04.08.487609.

3. Madeira F, Pearce M, Tivey ARN, Basutkar P, Lee J, Edbali O, Madhusoodanan N, Kolesnikov A, Lopez R. 2022. Search and sequence analysis tools services from EMBL-EBI in 2022. Nucleic Acids Res 50.

4. Letunic I, Bork P. 2021. Interactive tree of life (iTOL) v5: An online tool for phylogenetic tree display and annotation. Nucleic Acids Res 49.

5. Sievers F, Wilm A, Dineen D, Gibson TJ, Karplus K, Li W, Lopez R, McWilliam H, Remmert M, Söding J, Thompson JD, Higgins DG. 2011. Fast, scalable generation of high-quality protein multiple sequence alignments using Clustal Omega. Mol Syst Biol 7.

6. Anantharaman V, Makarova KS, Burroughs AM, Koonin E V., Aravind L. 2013. Comprehensive analysis of the HEPN superfamily: Identification of novel roles in intra-genomic conflicts, defense, pathogenesis and RNA processing. Biol Direct 8.

7. Dedrick RM, Jacobs-Sera D, Guerrero Bustamante CA, Garlena RA, Mavrich TN, Pope WH, Cervantes Reyes JC, Russell DA, Adair T, Alvey R, Bonilla JA, Bricker JS, Brown BR, Byrnes D, Cresawn SG, Davis WB, Dickson LA, Edgington NP, Findley AM, Golebiewska U, Grose JH, Hayes CF, Hughes LE, Hutchison KW, Isern S, Johnson AA, Kenna MA, Klyczek KK, Mageeney CM, Michael SF, Molloy SD, Montgomery MT, Neitzel J, Page ST, Pizzorno MC, Poxleitner MK, Rinehart CA, Robinson CJ, Rubin MR, Teyim JN, Vazquez E, Ware VC, Washington J, Hatfull GF. 2017. Prophage-mediated defence against viral attack and viral counter-defence. Nat Microbiol 2.

8. Mageeney CM, Mohammed HT, Dies M, Anbari S, Cudkevich N, Chen Y, Buceta J, Ware VC. 2020. *Mycobacterium* Phage Butters-Encoded Proteins Contribute to Host Defense against Viral Attack. mSystems 5.

